# Dysregulation of adipose ILC2 underlies thermogenic failure in aging

**DOI:** 10.1101/2020.09.08.288431

**Authors:** Emily L. Goldberg, Irina Shchukina, Yun-Hee Youm, Christina D. Camell, Tamara Dlugos, Maxim N. Artyomov, Vishwa Deep Dixit

## Abstract

Aging impairs the integrated immunometabolic responses which have evolved to maintain core body temperature in homeotherms to survive cold-stress, infections, and dietary restriction. Adipose tissue inflammation regulates the thermogenic stress response but how adipose tissue-resident cells instigate thermogenic failure in aged are unknown. Here, we define alterations in the adipose-resident immune system and identify that type 2 innate lymphoid cells (ILC2) are lost in aging. Restoration of ILC2 numbers in aged mice to levels seen in adults through IL-33 supplementation failed to rescue old mice from metabolic impairment and cold-induced lethality. Transcriptomic analyses revealed intrinsic defects in aged ILC2, and adoptive transfer of adult ILC2 are sufficient to protect old mice against cold. Thus, the functional defects in adipose ILC2 during aging drive thermogenic failure.

**One Sentence Summary:** Age-related changes in adipose tissue drive reprogramming of ILC2 that leads to impaired cold tolerance

## Main Text

Increasing evidence implicates the immune system as a key node in maintaining adipose tissue homeostasis by controlling the immunometabolic set point of inflammation. The adipose-resident immune compartment controls inflammation in visceral adipose tissue (VAT) and regulates fasting-induced lipolysis (*1, 2*), cold-induced thermogenesis (*3, 4*), and insulin-resistance (*5, 6*). While decades of research have pinpointed numerous defects in innate and adaptive arms of the anti-microbial immune response (*7–9*), only a limited number of studies have examined how tissue-resident immune cells that are responsible for maintaining tissue homeostasis regulate the aging process and resilience to stress (*10–12*).

Many of the physiological protective mechanisms described above decline with age, leading us to hypothesize that the adipose-resident immune compartment becomes dysfunctional during aging. To broadly characterize the VAT-resident immune compartments in adult and old mice we performed intravascular (iv) labeling to isolate exclusively tissue resident cells (*13*) followed by single-cell RNA sequencing (scRNAseq) (Figs. 1A-D, equal labeling efficiency demonstrated in Fig. S1, adult data was published previously (GSE137076) (*14*)). In agreement with prior studies, we observed losses in macrophages (clusters 0 and 4) (*15*), and increases in CD4 and CD8 αβ T cells (cluster 1) and regulatory T cells (Tregs, cluster 8) (*16*). We also identified profound losses of NK (cluster 5) and NKT cells (cluster 3) in aged adipose tissue. Amongst the most striking changes was an almost complete loss of type 2 innate lymphoid cells (ILC2, cluster 2) (Figs. 1B, C), which we verified by flow cytometry (Fig. 1E, F). We were especially intrigued by the loss of ILC2 in aged visceral adipose tissue because of their important role in promoting metabolic health and cold tolerance (*17–19*), both of which are impaired during aging. ILC2 exhibit strong tissue specificity (*20*) and are uniquely regulated in different tissues during the aging process (*21–23*), indicating that specific regulatory mechanisms might be responsible for their depletion in aged adipose tissue.

**Fig. 1.**
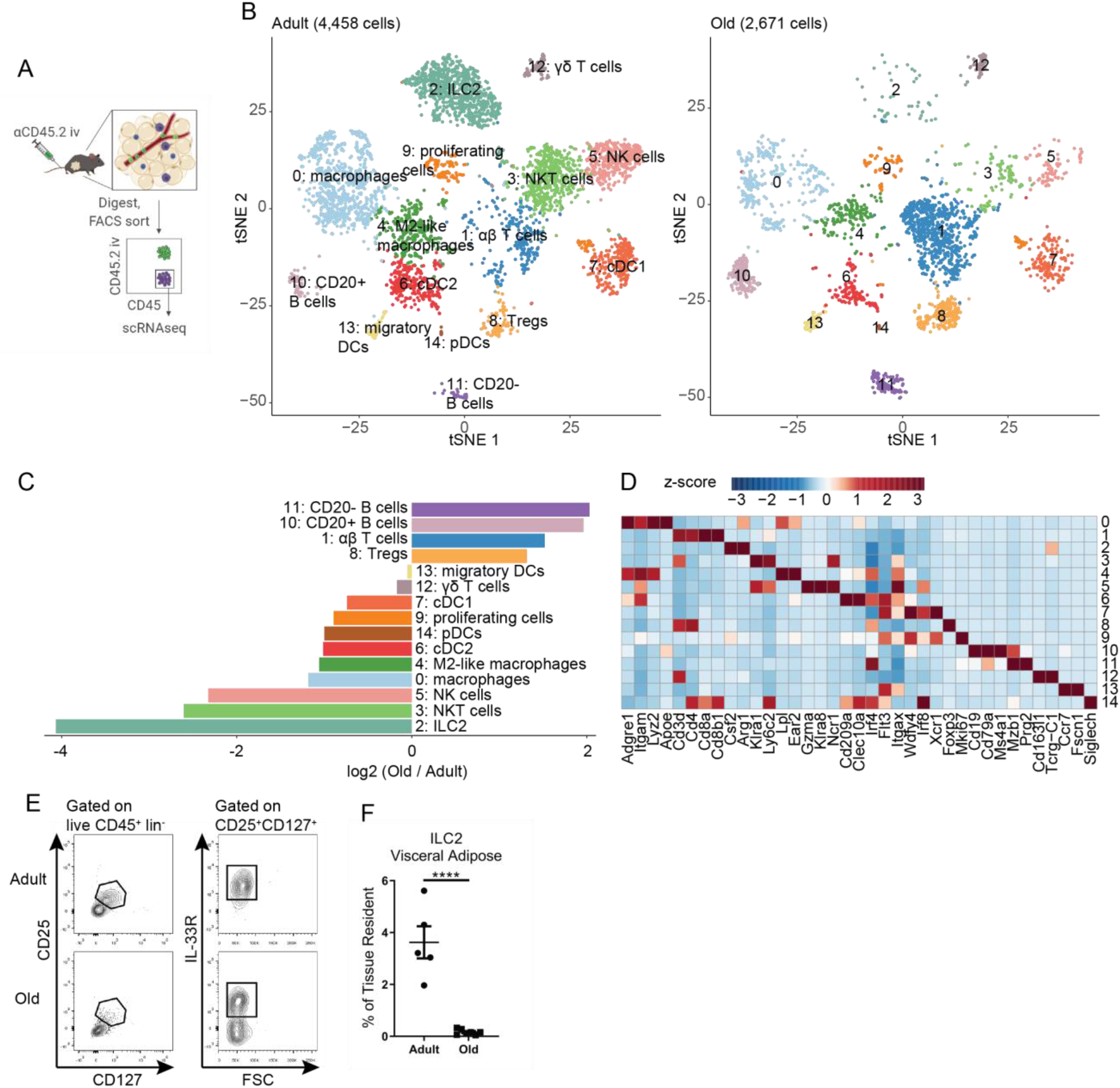
The adipose-resident immune compartment changes during aging. (A) Schematic depicting experimental strategy to define the adipose-resident immune compartment. (B) tSNE plots of tissue-resident CD45+ cells from adult (left) and old (right) male gonadal adipose tissue. (C) Bar chart showing population fold changes in relative abundance of each cluster induced by aging. (D) Heat map showing main lineage-defining genes to identify each cluster. (B-D) Data were generated from pooling n=4 adult and n=3 old male mice, so that a total of 1g of adipose tissue was used to isolate cells for each sample. (E) Representative flow cytometry analysis of gonadal adipose tissue ILC2 gating. (F) Quantification of ILC2 abundance in visceral adipose tissue in adult and old mice. (E-F) Data are representative of more than 10 experiments. Statistical differences in (F) were calculated by unpaired t-test.

ILC2 are seeded early in life and then retained within their respective tissues (*24*). Consistent with these data, we found that VAT ILC2 remain tissue-resident throughout lifespan (Figs. 2A, B) and that their loss occurred in both male and female mice (Fig. 2C). As ILC2 are an important source of IL-5 and therefore a primary regulator of eosinophils in visceral adipose tissue, we also found that eosinophils decline in both male and female mice during aging (Fig. 2D). Because obesity and age-related inflammation are associated with loss of ILC2 in adipose tissue (*19, 21*), we wondered whether the accumulation of fat mass might drive the ILC2 decline. We assessed the abundance of ILC2 in VAT in aged (15-month-old male) life-long calorie-restricted (CR) mice, which do not gain weight or fat mass during aging, and compared this to *ad libitum* fed age-matched controls, and young mice. Indeed, we found that the proportion of ILC2 (Fig. 2E), but not eosinophils (Fig. 2F), was retained in the VAT of aged CR mice. The actual number of ILC2 and eosinophils, normalized to tissue weight, were not preserved (Fig. S2) presumably because CR induces peripheral leukocytopenia (*25–28*). We have previously demonstrated that obesity and age-related adipose tissue inflammation leading to impaired metabolic health is driven by chronic low-level activation of the NLRP3 inflammasome (*29, 30*). However, aged *Nlrp3^-/-^* mice do not retain adipose ILC2 (Fig. 2G), indicating that their loss may be mediated by weight gain but is NLRP3-independent.

**Fig 2.**
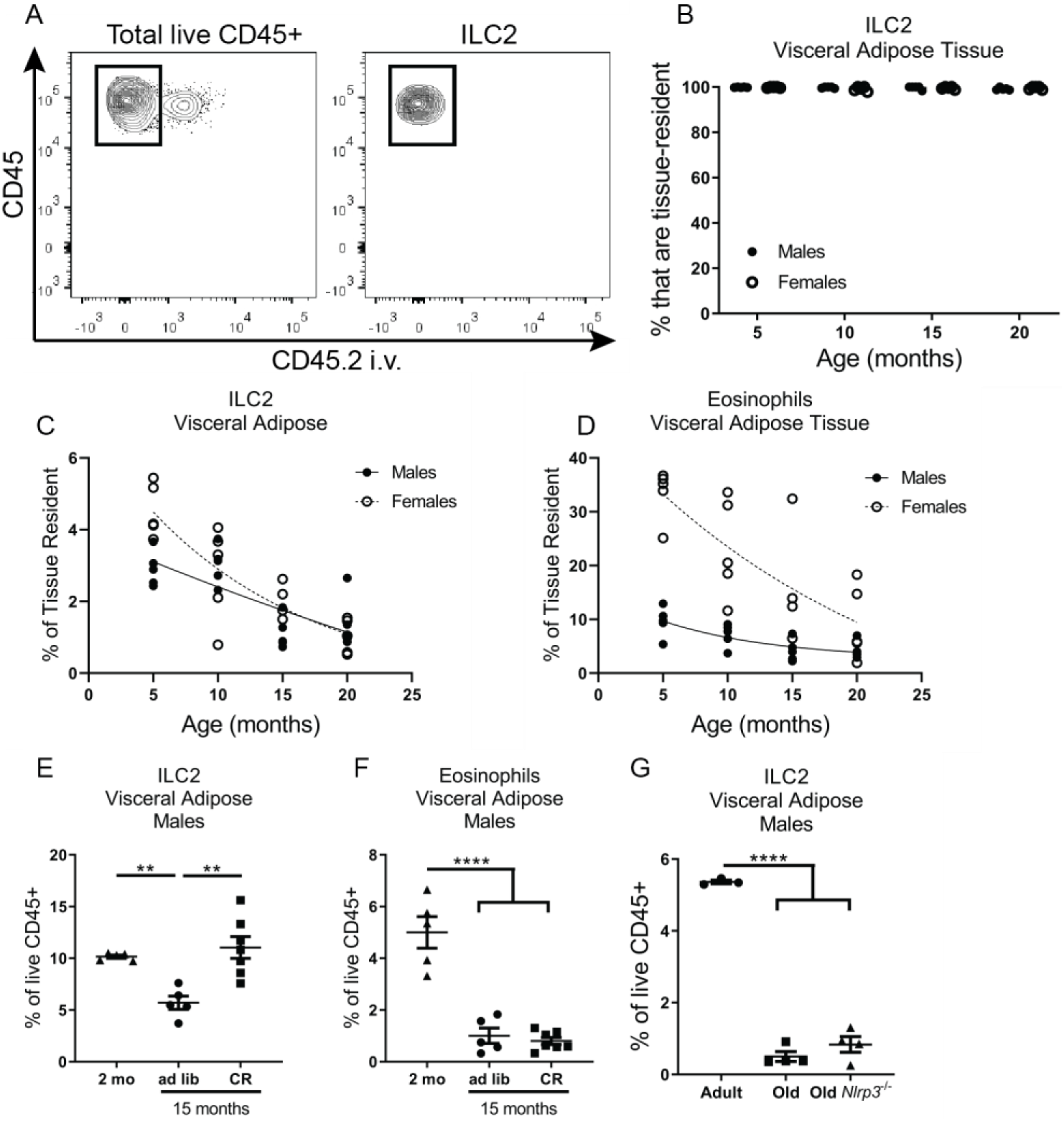
Loss of ILC2 during aging is sex-independent but driven by inflammation. (A) Representative gating scheme defining tissue-residence within different visceral adipose immune populations. (B) The percentage of ILC2 that are tissue resident (iv-label negative) in males and females across lifespan. The percentage of tissue-resident CD45+ cells that are (C) ILC2 and (D) eosinophils in aging male and female mice. Trend lines in (C, D) are best-fit lines for males and females, separately. For (A-D) data are representative of 5 independent experiments; n=5/group and each symbol represents an individual mouse. The percentage of adipose immune cells that are (E) ILC2 and (F) eosinophils in 15-month-old calorie-restricted (CR) male mice compared to adult and *ad libitum* age-matched control male mice. (G) Abundance of ILC2 in gonadal adipose tissue from adult, old, and age-matched old *Nlrp3^-/-^* male mice. For (E-G) Each symbol represents an individual mouse (n=4-7/group) and are representative of 2 independent experiments. Statistical differences were calculated by 1-way ANOVA with Tukey’s correction for multiple comparisons.

IL-33 is the primary survival cytokine that regulates ILC2 in VAT. IL-33 is produced by PDGFRα^+^ mesenchymal progenitors, and perivascular and endothelial cells in young mice (*3, 31–34*). However, when we examined which cell types express *Il33* in old mice using scRNAseq of the entire VAT stromal-vascular fraction, rather than detecting *Il33* expression in the *Pdgfra*^+^ cluster, we found that in aged mice nearly all *Il33* expression was restricted to a cluster of cells expressing the mesothelial cell marker *Msln* (Fig. 3A, Fig. S3). These data indicate that the cellular source of IL-33 in adipose tissue is switched during aging. Gene expression analysis by qPCR revealed an overall increase in *Il33* expression in VAT, but not subcutaneous adipose tissue, from old mice compared to young adult mice (Figs. 3B,C). These data suggest that alterations in cellular source of IL-33 may not allow ILC2s to access appropriate tonic signals required for maintenance in visceral adipose tissue during aging. Interestingly, concomitant with increase the in IL-33, aging led to specific increased expression of an *Il1rl1* transcript variant that encodes for soluble IL-33R in aged VAT with no change in serum levels (Figs. 3D-F). Given the increase in IL-33 expression, together with higher soluble IL-33R, it is likely that there is IL-33 signaling resistance in aged VAT that impairs ILC2 maintenance. However, ip injections of exogenous IL-33 in young and old mice caused equivalent STAT3 activation suggesting that IL-33 signaling is intact in aged adipose tissue. (Fig. 3G). Notably, the cellular source of soluble IL-33R is unknown in these studies. Additionally, whether increased soluble IL-33R is causal versus a consequence of elevated IL-33 in aged adipose remains to be determined. Regardless, these data provide a plausible mechanism by which IL-33 bioavailability may become reduced during aging, leading to loss of ILC2 in adipose tissue.

**Fig 3.**
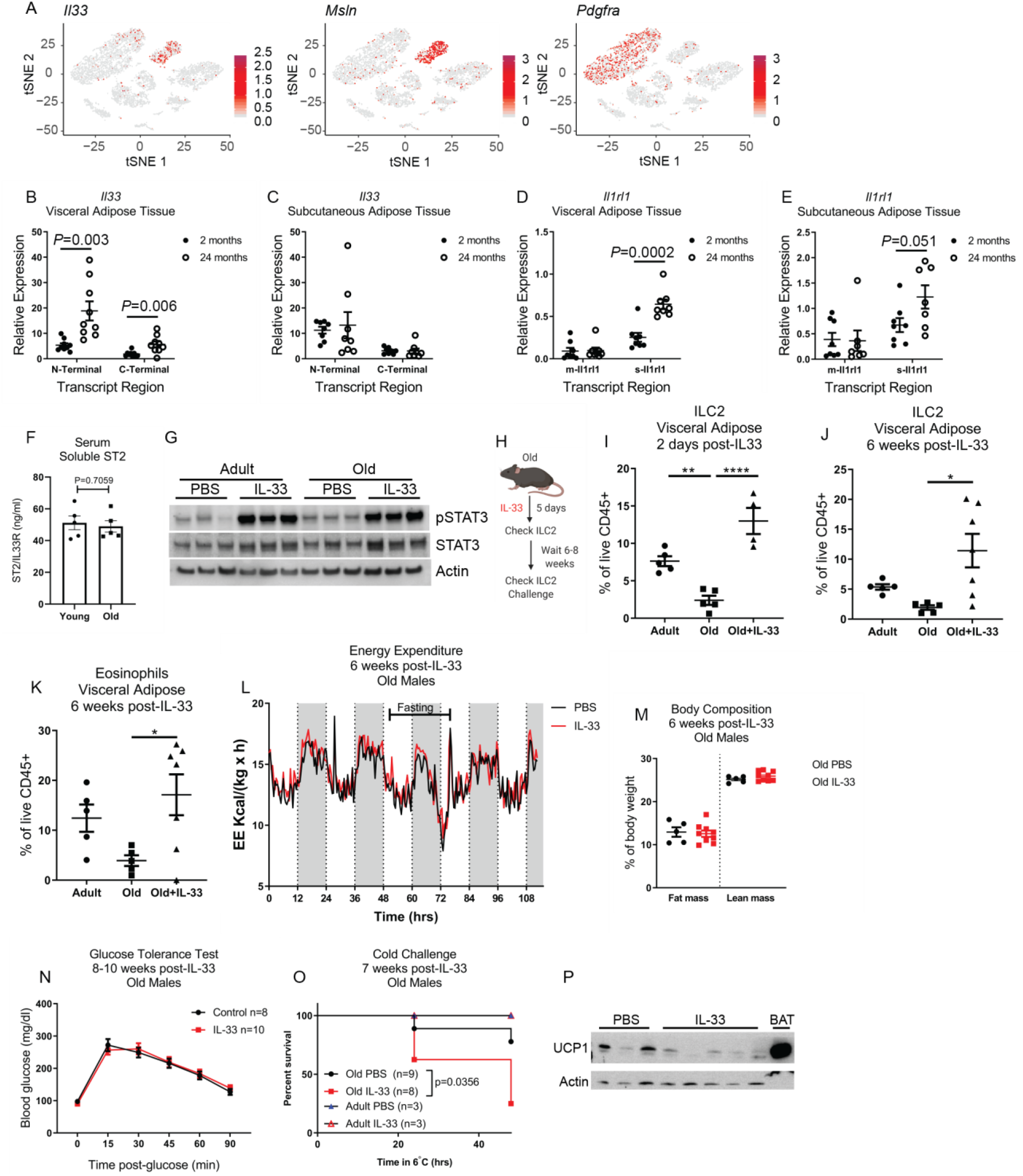
Aged ILC2 are IL-33-responsive but not metabolically protective. (A) tSNE plots of scRNAseq analysis of total SVF of gonadal adipose tissue obtained from a pooled samples of n=4 old female mice. Expression of *Il33, Msln*, and *Pdgfra*, are overlaid individually in red. For (B, C) *Il33* gene expression was measured by qPCR in (B) gonadal and (C) subcutaneous inguinal adipose tissue of adult and old mice. Primers were designed to measure relative expression of both N-terminal and C-terminal regions of *Il33*. Data are pooled from 2 independent experiments, for a total of n=9 VAT and n=8 SAT samples for each age group. Statistical differences between adult versus old mice were calculated by unpaired t-test for each transcript region analyzed. For (D, E) *Il1rl1* gene expression was measured in (D) gonadal and (E) subcutaneous inguinal adipose tissue of adult and old mice. Primers were designed to discriminate between the membrane-bound and soluble transcript variants. Data are pooled from 2 independent experiments, for a total of n=9mice/group for visceral adipose and n=8 mice/group for subcutaneous adipose samples. Each dot represents an individual mouse. Statistical differences between adult versus old mice were calculated by unpaired t-test for each transcript region analyzed. (F) Soluble IL33R (ST2) was measured in serum by ELISA. Statistical p-value was calculated by unpaired t-test. (G) Western blots of whole visceral adipose tissue lysate from adult and old mice treated with IL-33 or PBS. Each lane represents an individual mouse and data are representative of two independent experiments. (H) Schematic depicting experimental design. (I) ILC2 2 days after completing IL-33 treatments. (J) ILC2 and (K) eosinophils 6 weeks after completion of IL-33 treatment. Data are representative of 5 independent experiments, assessed through a range of 6-10 weeks after completing IL-33 treatments. Statistical differences were calculated by 1-way ANOVA with Tukey’s correction for multiple comparisons. (L) Energy expenditure in the fed and fasting state, measured 6 weeks after IL-33 treatment in old male mice. Data are representative of 2 independent cohorts. (M) Body composition in old male mice 6 weeks after IL-33 treatment. Data are representative of 2 independent experiments. (N) Glucose tolerance test in old male mice 8-10 weeks after IL-33 treatment. Data are pooled from 2 independent experiments. (O) Survival of male mice subjected to cold challenge at 7 weeks post-IL-33 treatment. Data are pooled from 2 independent experiments and were analyzed by log-rank test. (P) Protein expression of UCP1 in subcutaneous (inguinal) adipose tissue was measured by western blot in mice that survived 48hrs cold challenge. Data are representative of 2 independent experiments.

Next, we tested if providing excess IL-33 could override its apparent dysregulation to recover ILC2 in old mice (Fig. 3H). Consistent with the above data, treating old mice with IL-33 restored the numbers of ILC2 in aged adipose tissue (Fig. 3I). Co-administration of the sphingosine-1-phosphate receptor (S1PR) antagonist, FTY720, did not block ILC2 expansion, indicating this was due to *in situ* proliferation (Fig. S4A). However, given that IL-33 can be an alarmin in the IL-1 cytokine family and stimulates NFκB (*35*), and that immediately following IL-33 treatment there is also substantial eosinophilia in adipose tissue (Fig. S4B), we wondered whether the expanded ILC2 generates a sustained biological impact on aged VAT, which may also provide a longer therapeutic window for potential restorative effects of ILC2 on adipose tissue metabolism. Indeed, we measured increased numbers of ILC2 (Fig. 3J) and eosinophils (Fig. 3K) in aged VAT at least 6-8 weeks following completion of IL-33 treatment in aged mice.

We next considered the possibility that Tregs, which increase with age in male VAT and also expand in response to IL-33 (Fig. S5A-D), might outcompete ILC2 for cytokine survival factors because both subsets have uniquely high expression of *Il1rl1, Il7r*, and *Il2ra* (Fig. S5E-G) compared to all other adipose-resident immune cells. This hypothesis is also supported by previous studies showing that adipose Tregs disrupt metabolic health during aging (*36*), in contrast to their protective role in obesity (*6*). However, given that the visceral adipose Treg expansion is largely restricted to males (Fig. S5A) and that *in vivo* dose response experiments actually indicate ILC2 have higher IL-33 sensitivity compared to Tregs (Fig. S5D), we concluded this was not a likely explanation for the loss of ILC2 during aging.

Next, we investigated whether the restoration of ILC2 in aged mice, with levels comparable to adult mice, can result in metabolic therapeutic benefits. Surprisingly, as compared with PBS-treated control old mice, IL-33 treatment did not enhance energy expenditure (EE, Fig. 3L) or affect respiratory exchange ratio (RER, Fig. S6A) despite being reported to do so in young adult animals (*19*). Moreover, unlike what has been reported in young mice (*19*), IL-33 treatment also did not change fat mass/body composition (Fig. 3M) or glucose tolerance (Fig. 3N) in old mice. Additionally, consistent with above data that exogenous IL-33 doesn’t affect fatty acid substrate availability in aged VAT, the energetic parameters and induction of fasting-induced lipolysis were also not improved following IL-33 treatment (Fig. S6). However, we cannot rule out the possibility that the IL-33-induced expansion of Tregs in these mice might antagonize any metabolic benefits of the ILC2 during aging. Another reported function of adipose ILC2 is to protect core body temperature in response to cold challenge by inducing adaptive thermogenesis through beiging of white adipose tissue (*18, 19*). To our surprise, old mice with IL-33-mediated ILC2 restoration actually exhibited increased mortality in response to cold challenge (Fig. 3O). Lower expression of UCP1 in their subcutaneous white adipose tissue indicated this was due to an inability to induce adaptive thermogenesis and beiging (Fig. 3P).

Impaired thermogenic responses and increased mortality in IL-33-treated old mice after cold challenge raised the possibility that aged ILC2, and possibly the expanded eosinophils, were somehow defective. To test this possibility, we treated adult and old mice with IL-33 and then sorted visceral adipose tissue ILC2 for bulk RNAseq (Fig. 4A-C). Of note, we were unable to obtain reliable RNAseq data from sorted ILC2 from untreated old mice, likely due to how few ILC2 remain in aged fat. Gene Set Enrichment Analysis revealed significant alterations in transcriptional programs of ILC2 in adult and old mice (Fig. 4B, Fig. S7). Adult IL-33-expanded ILC2 showed enrichment of proliferation, transcription, and translation pathways, consistent with their mechanisms of self-renewal. In contrast, old IL-33-expanded ILC2 were enriched in cytokine signaling and activation pathways. We further examined the genes in the most enriched pathway, the JAK/STAT KEGG pathway (Fig. 4C) in old ILC2, and found enhanced expression of *Il6, Oslm*, and *Lif*, which all encode cytokines in the IL-6 family that are capable of inducing thermogenesis (*37, 38*), but are also associated with inflammatory disease and “inflammaging (*39*).” Similarly, old ILC2 had increased expression of *Il13* which is a common ILC2 cytokine associated with asthma, but can sustain alternatively activated macrophages associated with improved adipose tissue inflammation (*17*). However, macrophages within the adipose-resident compartment actually decline with age, and the proportion of pro-inflammatory macrophages increases (*2, 15*), suggesting that despite cytokine mRNA expression, old ILC2 function is impaired and this could further compound their numerical loss. Notably, this premise of functionality versus numbers was recently demonstrated in adipose tissue eosinophils (*40*). We also noted enhanced expression of *Ifngr1, Ifnar1*, and *Ifnar2* in aged ILC2 (Fig. 4C). Increased inflammation in adipose tissue, and specifically IFNγ, has been linked to loss of ILC2 in obese and aged adipose tissue (*21*). Similarly, increased type 1 interferon signatures during aging have been reported in several tissues (*30, 41, 42*). These data demonstrate that old ILC2 may be highly susceptible to cytokine changes in aging adipose tissue, and that old ILC2 might skew towards a hyperinflammatory phenotype with intrinsically altered survival mechanisms. Finally, to directly test if old ILC2 are dysfunctional, we performed adoptive transfer experiments. We sorted adult ILC2 from visceral adipose tissue of adult IL-5-RFP (Red5) reporter mice after IL-33-mediated expansion *in vivo* (Fig. 4D). Old mice that received adult ILC2 were protected against cold challenge (Fig. 4E) although they still exhibited some age-related impairment in maintaining their core body temperatures (Fig. 4F). Importantly, transferring adult ILC2 to old mice is sufficient to rescue susceptibility to cold challenge, indicating that aged ILC2 are intrinsically defective.

**Fig 4.**
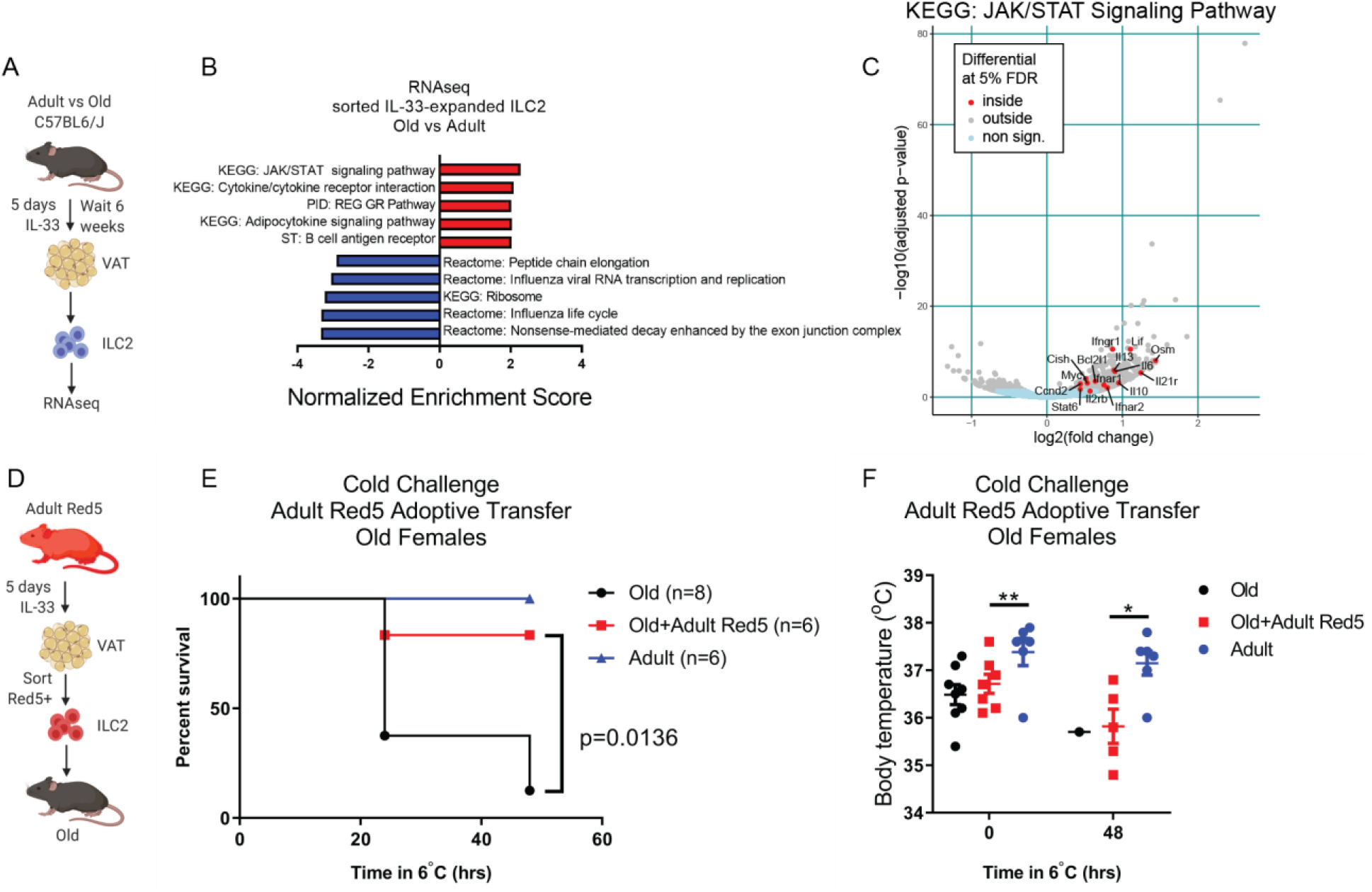
Intrinsic defects in aged ILC2 predispose to cold intolerance. (A) ILC2 bulk RNAseq experimental design. (B) Gene set enrichment analysis of IL-33-expanded ILC2 from adult and old mice. N=3 adult and n=3 old technical replicates, each containing sorted ILC2 pooled from n=2 biological replicate male mice were used for bulk RNAseq analyses. (C) Volcano plot highlighting significantly-regulated genes from the KEGG JAK/STAT Signaling Pathway gene list. (D) Adoptive transfer experimental design. (E) Survival and (F) body temperatures of old female mice during cold challenge ± receiving adult female Red5 ILC2 cells. Date are pooled from 2 independent experiments. Statistical differences in survival curves were analyzed by log-rank test. Statistical differences in body temperature were calculated by 2-way ANOVA.

In summary, our study identifies both cell-intrinsic and -extrinsic mechanisms leading to numerical and functional loss of ILC2 in aging adipose tissue. Given the important roles reported for ILC2 in mediating glucose homeostasis and inducing thermogenesis, we were surprised that restoring old ILC2 not only did not improve either of these outcomes in old mice, but instead impaired tissue homeostasis and increased susceptibility to cold challenge. It is unclear why a different source of IL-33 in aged adipose might detrimentally alter ILC2. Mesothelial cell-derived IL-33 has been identified as a rare source of IL-33 that acts as an alarmin to promote peritoneal inflammation after infection (*34*). But how mesothelial-versus mesenchymal-derived IL-33 could lead to ILC2 dysfunction requires further investigation. It is possible that the different locations of these cell types within the tissue might limit IL-33 availability to ILC2, whose localization is highly enriched in perivascular adventitia within adipose tissue (*31*). This likelihood is also supported by a recent study showing that cell-cell interactions between ILC2 and IL-33-producing PDGFRα^+^ cells are important for ILC2 maintenance/proliferation (*43*). Similarly, increased gene expression of soluble IL33R could reduce bioavailability of IL-33 to adipose-resident ILC2. Collectively, these data support a model in which loss of IL-33 availability drives the age-related loss in ILC2. Moreover, ILC2 might adapt to this limitation, and become permanently reprogrammed in the process. In this case, when expanded with IL-33, a pool of pathogenic ILC2s multiply in aged adipose that are no longer capable of normal ILC2 functions. Alternatively, due to their acquisition of proinflammatory cytokine production, it is possible that the loss of ILC2s during aging reflects a programmed adaptive protective mechanism. In conclusion, these data highlight the importance of studying changes in tissue-resident immune compartments to gain deeper understanding of the mechanisms contributing to tissue functional decline and loss of homeostasis during aging.

## Materials and Methods

### Mice

All mice were on the C57BL/6J genetic background. Adult (8-12 weeks old) and old (18-24 months old) mice were obtained from Jackson Labs or the National Institute on Aging (NIA) Aged Rodent Colony, and comparisons were made between groups of mice from the same vendor. Red5 mice were purchased from Jackson Labs (stock #030926) and were originally generated and described by Nussbaum et al (*44*). *Nlrp3^-/-^* mice have been described previously (*45*) and were bred and aged in the Dixit lab, as were the control wildtype mice. Calorie-restricted mice and their age-matched *ad libitum* controls were obtained from the NIA Aged Rodent Colony. Rag^-/-^IL-2Rγ^-/-^ mice were purchased from Taconic (model #4111). Mice were housed under standard 12hr light/dark cycles. All animal procedures were approved by the Yale Institutional Animal Care and Use Committee.

### IL-33 Treatments

Mice were treated by i.p. injection of IL-33 (Biolegend) at a dose of 12.5ug/kg body weight, unless a different dose is specifically indicated in the figure, for 5 consecutive days. Mice were analyzed early (2 days) or late (at least 6-8 weeks) after completing IL-33 treatments. Where indicated, mice were co-treated with FTY-720 (1mg/kg) every other day.

### Glucose Tolerance Tests

Mice were fasted 16hr prior to glucose tolerance test. Glucose was given by i.p. injection based on body weight (0.4g/kg) and blood glucose levels were measured by handheld glucometer.

### Body Composition and Energy Expenditure

Body composition was measured *in vivo* by magnetic resonance imaging (EchoMRI; Echo Medical Systems). The TSE PhenoMaster System (V3.0.3) Indirect Calorimetry System was used to monitor energy expenditure, activity, and food and water intake in individual mice at 3Omin intervals. Oxygen consumption and carbon dioxide production measurements were used to determine energy expenditure.

### IL33R (ST2) Measurements

IL33R protein levels were measured in serum by ELISA (R&D) according to the manufacturer’s instructions. Relative expression of soluble and membrane-bound IL33R expression was measured by qPCR. Primer sequences are provided in Supplemental Table 1.

### Western blots

Mice were treated with IL-33 for 5 consecutive days, as described above. 30 minutes after their final IL-33 injection, visceral adipose tissue was collected and snap frozen in liquid nitrogen. Tissue was homogenized in RIPA buffer containing protease inhibitors (Pierce) and protein expression was assessed by western blot. Antibodies against phosphorylated STAT3 (pSTAT3, 1:1000, Tyr705 rabbit, Cell signaling), total STAT3 (1:1000; 79D7 rabbit, Cell Signaling), and β-actin (1:1,000 4967L; Cell Signaling) and HRP-conjugated goat-anti-rabbit secondary antibody (PI31460; Thermo Scientific) were used to visualize proteins using chemiluminescence (PI32106; Thermo Scientific).

### Flow Cytometry

Intravascular labeling was performed by iv injection of 2.5ug CD45.2-FITC diluted in 100ul PBS. Mice were euthanized exactly 3 minutes after injection for tissue collection. Adipose tissue was digested in HBSS + 1mg/mL Collagenase I in shaking 37°C water bath. Cells were stained with live/dead viability dye (Invitrogen), incubated with Fc block, and then stained for surface markers including CD45, CD127, CD25, IL-33R (ST2), and a dump channel that included CD3, CD19, γδ TCR, CD4, CD8, NK1.1, CD11b, F4/80, Ly6G, Ly6C, and Ter-119. All antibodies were purchased from eBioscience or Biolegend. When needed, intracellular staining for Foxp3 was performed using the eBioscience Fix/Perm nuclear staining kit, otherwise cells were fixed in 2% PFA. Samples were acquired on a custom LSR II and data was analyzed in FlowJo.

### In vitro lipolysis assay

Glycerol release from adipose tissue explants was used to asses induction of lipolysis. Approximately 10mg of adipose was used for each assay. Where indicated, 10^3^ sorted adult Red5 cells were added for the entire duration of the assay. Isoproterenol was added at a concentration of 1μM where indicated. Glycerol release was measured according to the manufacturer’s instructions (Sigma, MAK117).

### Single-cell RNA sequencing sample preparation

After iv-labeling epidydimal white adipose tissue was collected, digested, and stained for viability (live/dead Aqua, ThermoFisher) and pan-CD45 to FACS sort live tissue-resident hematopoietic cells. Adipose tissue from n=4 adult and n=3 old mice were pooled so that 1 gram of tissue for each age group was used for isolating tissue-resident CD45+ cells for sequencing. For total SVF analyses, visceral adipose tissue from n=4 old female mice were pooled. Cells were prepared for single-cell sequencing according to the 10X Genomics protocols. Sequencing was performed on a HiSeq4000.

### scRNAseq Alignment, barcode assignment and unique molecular identifier (UMI) counting

The Cell Ranger Single-Cell Software Suite (v2.1.1) (available at https://support.10xgenomics.com/single-cell-gene-expression/software/pipelines/latest/what-is-cell-ranger) was used to perform sample demultiplexing, barcode processing, and single-cell 3’ counting. Cellranger mkfastq was used to demultiplex raw base call files from the HiSeq4000 sequencer into sample-specific fastq files. Subsequently, fastq files for each sample were processed with cellranger counts to align reads to the mouse reference (version mm10-2.1.0).

The default estimated cell count value of 10,000 was used for this experiment. Samples were subsampled to have equal numbers of confidently mapped reads per cell.

### scRNAseq Preprocessing analysis with Seurat package

For the analysis, the R (v3.4.2) package Seurat (v2.3) (*46, 47*) was used. Cell Ranger filtered genes by barcode expression matrices were used as analysis inputs. Samples were pooled together using the AddSample function. The fraction of mitochondrial genes was calculated for every cell, and cells with high (>5%) mitochondrial fraction were filtered out. Expression measurements for each cell were normalized by total expression and then scaled to 10,000, after that log normalization was performed (NormalizeData function). Two sources of unwanted variation: UMI counts and fraction of mitochondrial reads - were removed with ScaleData function.

### scRNAseq Dimensionality reduction and clustering

The most variable genes were detected using the FindVariableGenes function. PCA was run only using these genes. Cells are represented with t-SNE (t-distributed Stochastic Neighbor Embedding) plots. We applied RunTSNE function to normalized data, using first 10 PCA components. For clustering, we used function FindClusters that implements SNN (shared nearest neighbor) modularity optimization-based clustering algorithm on top 10 PCA components using resolution of 0.5.

### scRNAseq Identification of cluster-specific genes and marker-based classification

To identify marker genes, FindAllMarkers function was used with likelihood-ratio test for single cell gene expression. For each cluster, only genes that were expressed in more than 10% of cells with at least 0.1-fold difference (log-scale) were considered. For heatmap representation, mean expression of markers inside each cluster was used.

### scRNAseq differential expression

To obtain differential expression between clusters, MAST test was performed, and p-value adjustment was done using the Bonferroni correction (*48*). Only genes that were expressed in more than 10% of cells in cluster were considered.

### Bulk RNA Isolation and Transcriptome Analysis

ILC2 were FACS sorted from epidydimal fat and RNA was isolated using the RLT method. RNA was sequenced on a HiSeq2500. Fastq files for each sample were aligned to the mm10 genome (Gencode, release M23) using STAR (v2.7.3a) with the following parameters: STAR --genomeDir $GENOME_DIR --readFilesIn $WORK_DIR/$FILE_1 $WORK_DIR/$FILE_2 --runThreadN 12 --readFilesCommand zcat --outFilterMultimapNmax 15 --outFilterMismatchNmax 6 -- outReadsUnmapped Fastx --outSAMstrandField intronMotif --outSAMtype BAM SortedByCoordinate --outFileNamePrefix./$ (*49*). Quality control was performed by FastQC (v0.11.3), MultiQC (v1.1) (*50*), and Picard tools (v2.18.4). Quantification was done using htseq-count function from HTSeq framework (v0.9.1): htseq-count -f bam -r pos -s no -t exon $BAM $ANNOTATION > $OUTPUT (*51*). Differential expression analysis was done using DESeq function from DeSeq2 package (*52*) (v1.24.0) with default settings. Significance threshold was set to adjusted p-value < 0.05. Gene set enrichment analysis via fgsea R package (*53*) (v1.10.0) was used to identify enriched pathways and plot enrichment curves.

### Adoptive Transfer

Adult Red5 mice were treated with IL-33 for 5 consecutive days as described above, rested for 2 days, and then RFP+ cells were sorted from digested visceral adipose tissue using a BD FACSAria. A total of 3×10^5^ sorted ILC2 were transferred into old sex-matched recipients in bilateral i.p. injections (1.5×10^5^ per injection), and recipient mice were simultaneously given a single dose of IL-33 at 12.5ug/kg body weight to facilitate engraftment. Recipient mice were rested for 1-2 weeks and then subjected to cold challenge.

### Cold Challenge

Mice were singly-housed in cages that contained bedding, but without nestlet, for cold challenge at 6°C. Mice were monitored twice daily for survival during cold challenge, and adipose tissue was collected from surviving mice at 48hr time point for western blot of UCP1 (abcam) expression.

### Statistical Analysis

All statistical tests for each experiment are indicated in the corresponding figure legend. For all experiments, p<0.05 was considered significant. *p<0.05, **p<0.01, ***p<0.001, ****p<0.0001.

## Acknowledgments

We thank all members of the Dixit lab for critical discussion and feedback related to this project. All schematics were created with BioRender.com

## Author contributions

EG and VDD conceived the overall project, experiments, and data interpretation. EG planned and performed all experiments, analyzed data, and prepared manuscript. IS performed all RNA sequencing analyses and provide the corresponding figure panels. YY performed aged SVF ssRNAseq experiment, and qPCR analyses. CDC assisted in experiments analyzing adipose-resident immune changes throughout aging. TD helped perform experiments involving IL-33 injections. MNA oversaw all RNAseq analyses. All authors assisted in manuscript preparation.

## Data and materials availability

All sequencing data will be deposited in the Gene Expression Omnibus (GEO) database and will be made publicly available upon publication of the paper.

