## Supplemental Figures for "Dysregulation of adipose ILC2 underlies thermogenic failure in aging"

**This PDF file includes:**

Figs. S1 to S7
Table S1

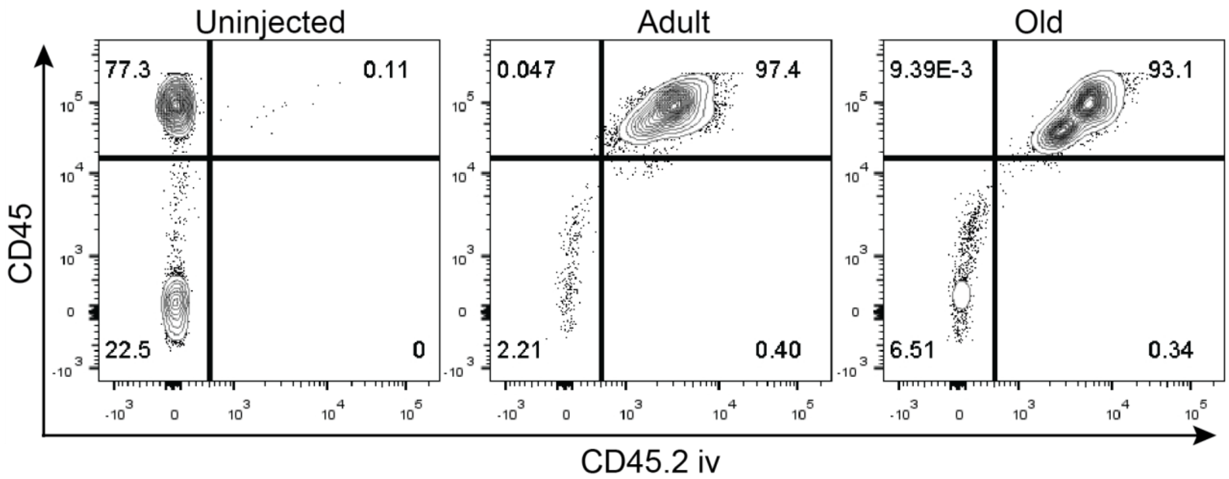


**Fig. S1.**
Flow cytometry staining of blood collected from adult and old mice after iv labeling.

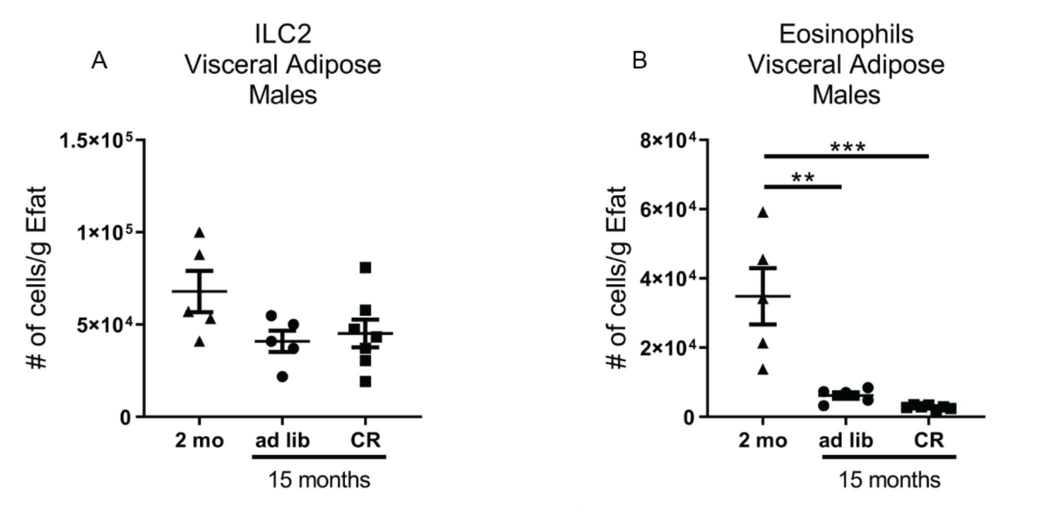


**Fig. S2.**

The numbers of (A) ILC2 and (B) eosinophils in gonadal adipose tissue of male mice were normalized to fat pad mass. Data are representative of 2 independent experiments. Each symbol represents an individual mouse. Statistical differences were calculated by 1-way ANOVA with Tukey's correction for multiple comparisons.

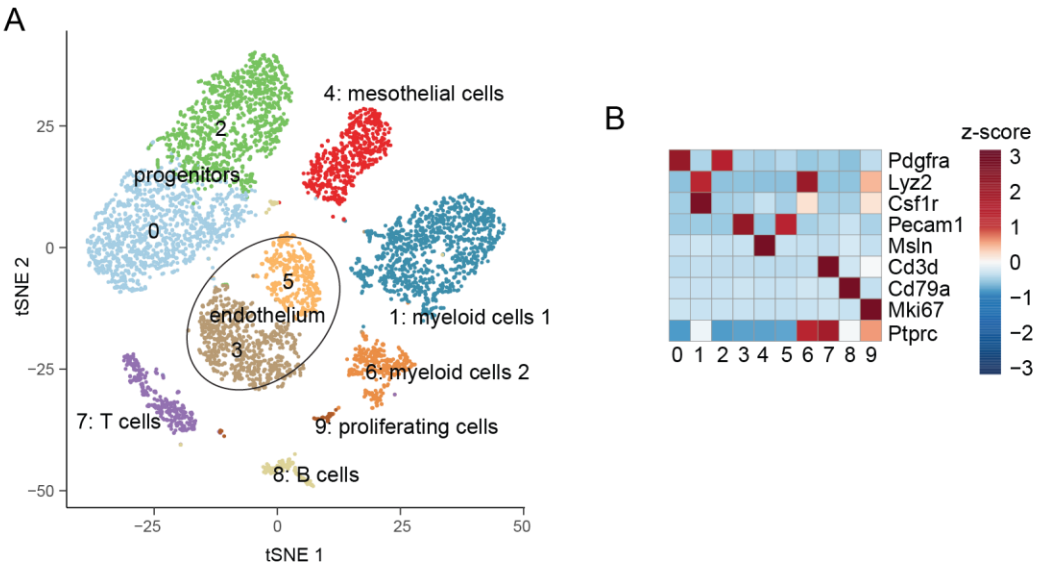


**Fig. S3.**

(A) tSNE plot of old female adipose tissue SVF scRNAseq. (B) Heat map showing main lineage-defining genes used to identify each cluster.

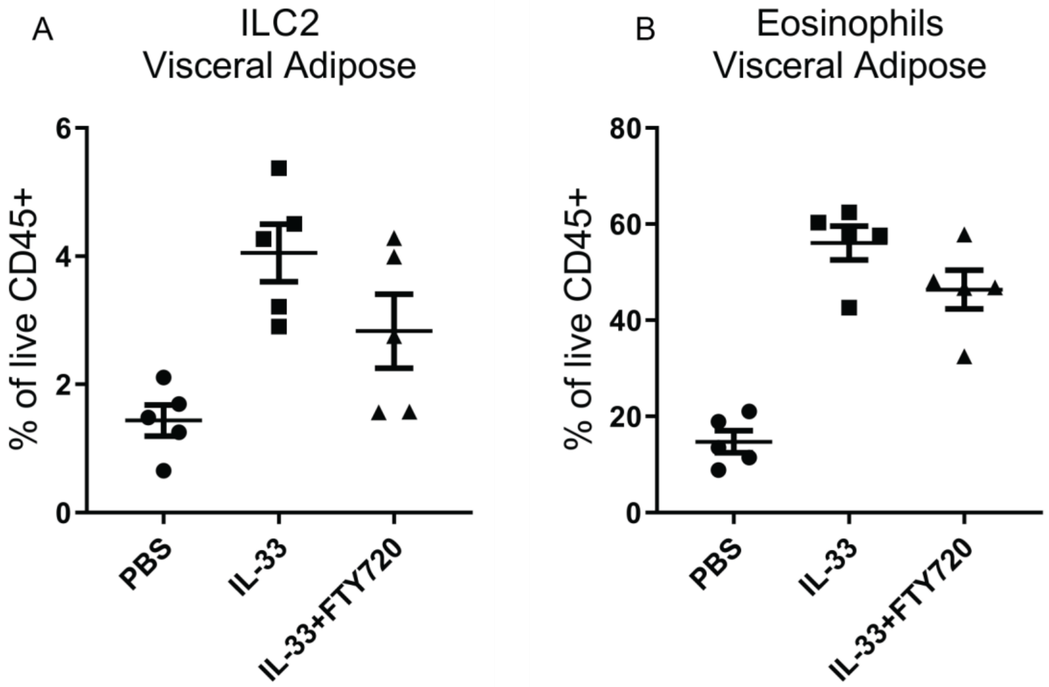


**Fig. S4.**
Old mice with treated with IL-33 ± FTY720 for one week. The following week, two days after completing IL-33 treatment, abundances of (A) ILC2 and (B) eosinophils in gonadal adipose tissue were assessed by flow cytometry. Each symbol represents an individual mouse. Data are representative of 2 independent experiments, each containing n=5 mice/group.

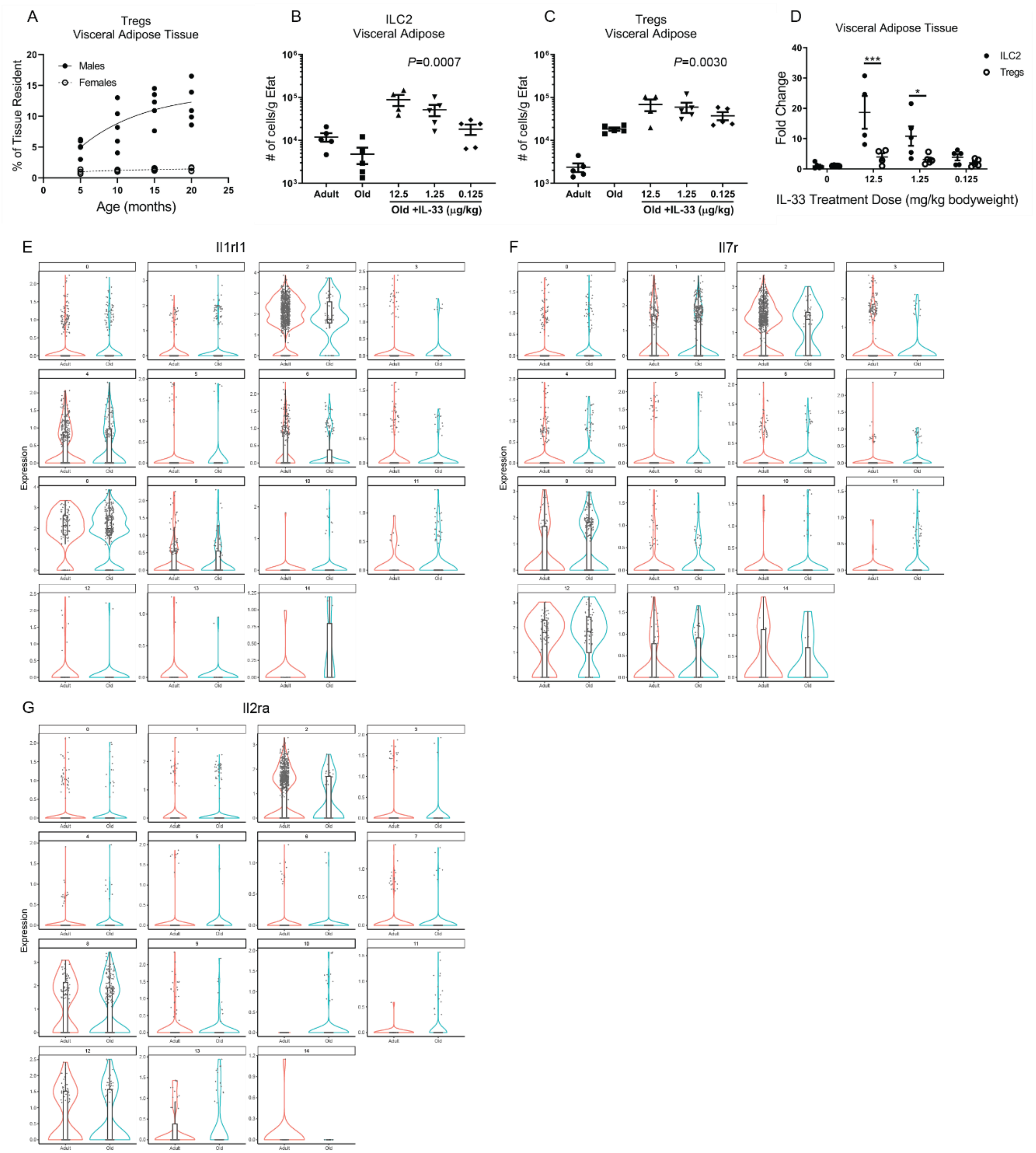


**Fig. S5.**
(A) Proportion of adipose-resident CD45+ cells that are Tregs in aging male and female mice. Data are representative of 3 independent experiments, each symbol represents an individual mouse. N=5 male and n=5 female per age group, trend line shows the "best fit." Numbers of (B) ILC2 and (C) Tregs in gonadal adipose tissue after 5 days of IL-33 treatment at the indicated

doses. Statistical differences were calculated by 1-way ANOVA. (D) Fold change of numbers of ILC2 and Tregs induced by IL-33 treatment in old mice. Statistical differences were calculated by 2-way ANOVA. Violin plots of (E) *Il1rl1*, (F) *Il7r*, and (G) *Il2ra* within each cluster of adult (red) and old (blue) adipose-resident CD45+ cells from Figure 1. For violin box plots, the middle line represents the median, and upper and lower hinges represent the 7 $5^{th}$ and $25^{th}$ percentiles, respectively. For (B-D) Mice were analyzed two days after completing IL-33 treatment. Each symbol represents an individual mouse and data are representative of 2 independent experiments.

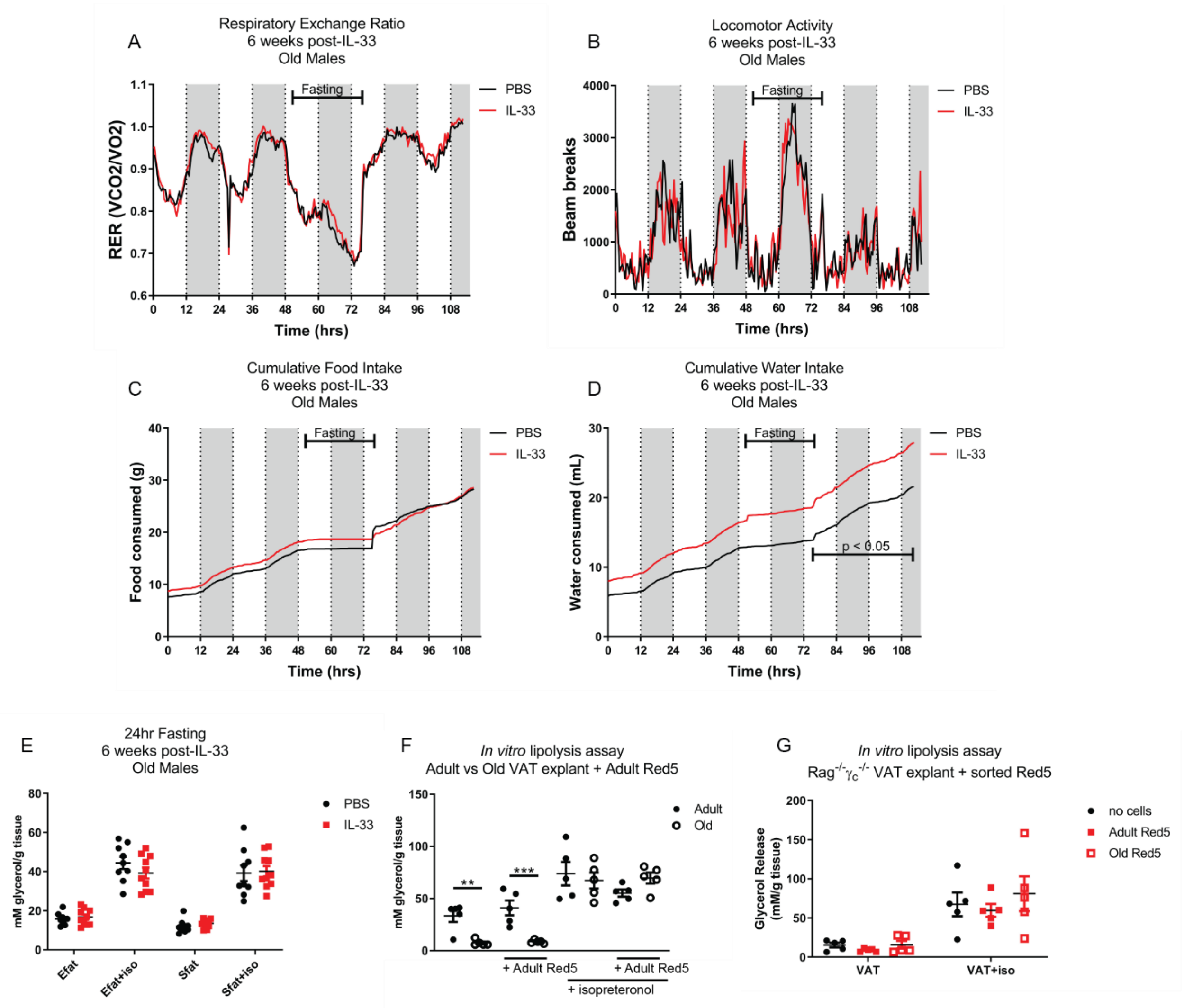


**Fig. S6.**
6 weeks after IL-33 treatment mice were housed in metabolic cages to measure (A) Respiratory exchange ratio (RER), (B) locomotor activity, (C) food intake, and (D) water intake. Data are representative of 2 independent experiments and statistical differences were assessed by paired 2-way ANOVA. (E) Glycerol release from visceral adipose tissue explants after 24hr fasting in old male mice that had been treated with PBS or IL-33 6 weeks earlier. Data are representative of 3 independent experiments. N=9 mice/group (PBS vs IL-33) and the same mouse was used to measure both Efat and Sfat ± isopreteronol (iso). (F) Glycerol release from adipose tissue explants obtained from fed adult and old mice. Explants were co-incubated with Adult Red5 ILC2 as indicated, and lipolysis was stimulated by addition of isopreteronol to the indicated sample groups. Each dot represents a unique adult (n=5) or old (n=5) source of adipose tissue, and tissue from the same mice were used for each condition. Statistical differences were calculated by 2-way ANOVA separately for the unstimulated and isopreteronol-stimulated samples. (G) Glycerol release from adipose tissue explants obtained from fed $Rag^{-/-}\gamma_c^{-/-}$ mice lacking ILC2. Explants were co-incubated with sorted Adult vs Old Red5 ILC2 as indicated, and lipolysis was stimulated by addition of iso to the indicated samples. Each dot represents a unique adult $Rag^{-/-}\gamma_c^{-/-}$ mouse (n=5), and tissue from the same mice were used for each condition. All figures are representative of at least 3 independent experiments.

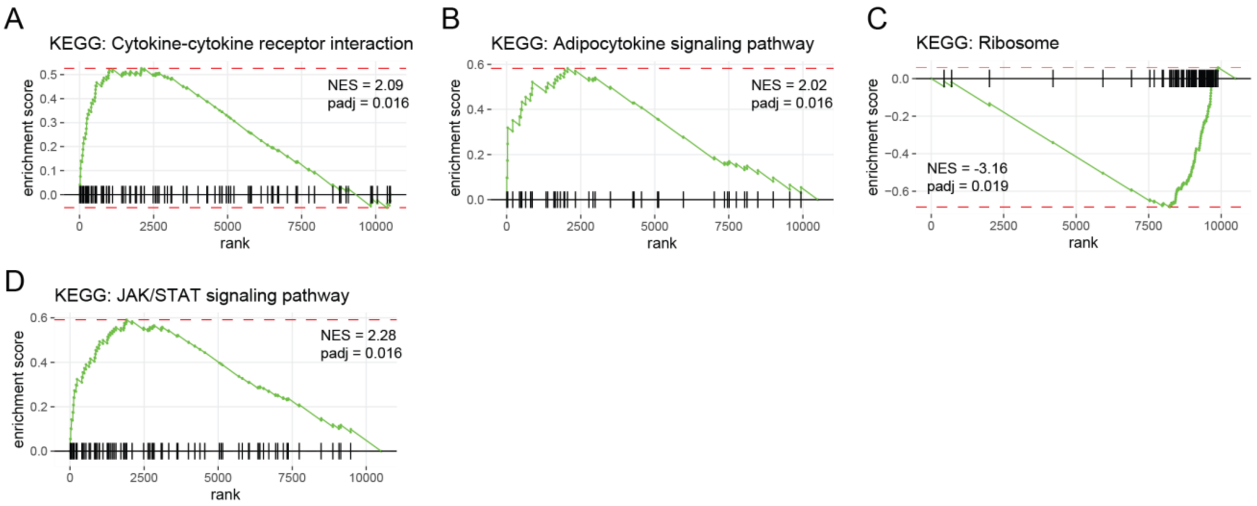


**Fig. S7.**
(A-D) GSEA enrichment curves of significantly regulated pathways identified in Figure 4B.

| Target | Template ID | Forward Primer (5’$\rightarrow$3’) | Forward Primer (5’$\rightarrow$3’) |
|---|---|---|---|
| Soluble IL33R (Variant 3) | NM_001294171.1 | TGGCTAGGACCTCTGGCTAA | ATGGTGTGTTCACTAGGCGG |
| Membrane-bound IL33R (Variant 1) | NC_000067.6 NM_001025602.3 Range: 40447012 - 40465336 | CTCTGCCCGACGTTCTTGA | AACCCCTGATGTGTCTCAGT |
| IL-33 N-terminal | NM_001164724.2 | AACTCCAAGATTTCCCCGGC | TTATGGTGAGGCCAGAACGG |
| IL-33 C-terminal | NM_001164724.2 | AGACCAGGTGCTACTACGCT | ACGTCACCCCTTTGAAGCTC |

**Table S1.**
Primer sequences used for qPCR gene expression analyses.
